# Regulation of nuclear-cytoplasmic partitioning by the *lin-28-lin-46* pathway reinforces microRNA repression of HBL-1 to confer robust cell-fate progression in *C. elegans*

**DOI:** 10.1101/698977

**Authors:** Orkan Ilbay, Victor Ambros

## Abstract

MicroRNAs target complementary mRNAs for degradation or translational repression, reducing or preventing protein synthesis. In *C. elegans*, the transcription factor HBL-1 (Hunchback-like 1) promotes early larval (L2) stage cell-fate, and the *let-7*-family microRNAs temporally down-regulate HBL-1 to enable the L2-to-L3 cell-fate progression. In parallel to *let-7*-family microRNAs, the conserved RNA binding protein LIN-28 and its downstream gene *lin-46*, also act upstream of HBL-1 in regulating the L2-to-L3 cell-fate progression. The molecular function of LIN-46, and how the *lin-28-lin-46* pathway regulates HBL-1, are not understood. Here, we report that the regulation of HBL-1 by the *lin-28-lin-46* pathway is independent of the *let-7/lin-4* microRNA complementary sites (LCSs) in the *hbl-1* 3’UTR, and involves a stage-specific post-translational regulation of HBL-1 nuclear accumulation. We find that LIN-46 is necessary and sufficient to prevent nuclear accumulation of HBL-1. Our results illuminate that the robust progression from L2 to L3 cell-fates depends on the combination of two distinct modes of HBL-1 down-regulation: decreased synthesis of HBL-1 via *let-7*-family microRNA activity, and decreased nuclear accumulation of HBL-1 via action of the *lin-28-lin-46* pathway. Like HBL-1, many microRNA targets are transcription factors (TFs); and cooperation between regulation of nuclear accumulation and microRNA-mediated control of synthesis rate may be required to increase the precision of or confer robustness to down-regulation of these microRNA target TFs, which can be critical to achieve the optimal phenotypes.

## Introduction

Precise and robust gene regulation is crucial for animal development. Optimal doses of developmental gene products expressed with spatiotemporal precision produce the wild-type body plan; whereas abnormally lower or higher doses or ectopic expression of developmental genes can result in morphological defects that reduce the fitness of the individual. The proper spatiotemporal activity of developmental gene products is ensured by elaborate gene regulatory mechanisms, which often involve collaboration across semi-redundant mechanisms controlling the gene activity at different levels – transcriptional, translational, and post-translational.

*Caenorhabditis elegans* (*C. elegans*) development consists of an invariant set of cell division and differentiation events that produces the stereotyped adult body plan [1]. *C. elegans* developmental regulators are identified by loss-of-function or gain-of-function mutations that cause developmental lethality, or evident morphological defects. One class of developmental defects in *C. elegans* stems from changes in the order and/or timing of larval developmental events, controlled by the heterochronic gene pathway [2]. In this pathway, three transcription factors (LIN-14, HBL-1, LIN-29) control cell-fate transitions from earlier to later stages, and are temporally regulated – directly or indirectly – by certain microRNAs. In particular, LIN-14 is regulated by *lin-4* [3], HBL-1 (Hunchback-like-1) is regulated by three members of the *let-7* family (a.k.a. the *let-7* sisters: *mir-48, mir-84* and *mir-241*) [4], and LIN-29 is post-transcriptionally regulated by the RNA binding protein LIN-41 [5], which is in turn regulated by *let-7* [6]. These microRNAs are dynamically expressed during larval development and they ensure proper temporal down-regulation of their targets, which is crucial for the proper program of stage-appropriate cell-fate transitions.

The conserved RNA binding protein LIN-28 also plays key roles in the *C. elegans* heterochronic pathway. *lin-28* regulates early cell-fates upstream of *lin-46* [7] and in parallel to *mir-48/84/241* [4], and regulates late cell-fates upstream of the conserved microRNA *let-7* [8,9]. In larvae lacking *lin-28*, hypodermal stem cells (called seam cells) skip L2 stage proliferative cell-fates and precociously express terminally differentiated adult cell-fates [2,10] while the rest of the tissues, e.g. the gonad, are still juvenile and developing. *lin-46(lf)* suppresses these *lin-28(lf)* phenotypes [7]: *lin-28(lf)*;*lin-46(lf)* double mutants are wild-type for the phenotypes observed in *lin-28(lf)* animals*. lin-28(lf)* also suppresses the heterochronic phenotypes of *mir-48/84/241* mutants [4], which is characterized as reiterations of the proliferative L2 cell-fates at later stages; and *lin-46* is also required for this suppression [11].

Animals lacking *lin-46* display weak heterochronic phenotypes that are enhanced when the larvae are cultured at low temperatures, such as 15°C, which otherwise do not affect wildtype larval development [7]. While wildtype larval development is similarly robust against other stresses such as population density pheromones or starvation [12], *lin-46(lf)* phenotypes are enhanced under these conditions [12], and when animals develop through traversing a temporary diapause in response to these stresses [13]. Interestingly, diapause-inducing conditions also repress *let-7*-family microRNA expression [14,15]; which is thought to be important for (optimal) diapause decision [15] and prolonging HBL-1 expression in coordination with the rate of developmental stage progression [12]. Therefore, LIN-46 activity is important for the proper down-regulation of HBL-1 under physiological conditions where *let-7*-family microRNA levels are reduced.

The molecular functions of LIN-46, and how these functions may relate to its role in the heterochronic pathway are not known. However, given the prominent involvement of microRNAs in the heterochronic pathway, in the context of regulating temporal cell-fates, LIN-46 has been thought to possibly act by modulating (more precisely “boosting”) the activities of certain microRNAs, e.g. *lin-4* or *let-7-*family microRNAs, perhaps by interacting with the miRISC (microRNA-induced silencing complex). By hypothetically boosting the activities of *lin-4* and/or *let-7*-family microRNAs in response to environmental stress, LIN-46 could compensate for the reduced levels of these microRNAs. This model is supported by the correlated conservation of LIN-46 with the argonaute-family proteins, suggesting a potential function for LIN-46 related to microRNA or small RNA pathways [16]. Alternatively, LIN-46 could regulate HBL-1 activity by a mechanism independent of microRNAs.

Here we report that deleting a genomic region encompassing the *let-7/lin-4* complementary sites (LCSs) in the *hbl-1* 3’UTR results in strong extra seam cell phenotypes, which is consistent with lack of *let-7/lin-4* microRNA regulation and a gain-of-function of HBL-1. Importantly, we find that *lin-28(lf)* suppresses, and *lin-46(lf)* enhances, the extra seam cell phenotype of *hbl-1(gf*/Δ*LCSs)*, indicating that the regulation of HBL-1 by the *lin-28-lin-46* pathway is independent of the LCSs. Moreover, HBL-1, which normally localizes to the nucleus, accumulates in the cytoplasm of hypodermal seam cells in *lin-28(lf)* and *hbl-1(gf*/Δ*LCSs)* animals, and *lin-46* is required for this cytoplasmic accumulation of HBL-1. This cytoplasmic accumulation is accompanied by reduced nuclear accumulation of HBL-1, which also correlates with reduced HBL-1 activity. Lastly, we found that precocious expression of LIN-46 in L2 stage seam cells is sufficient to localize HBL-1 to the cytoplasm, reducing the nuclear accumulation of HBL-1, and thereby suppressing *hbl-1* gain-of-function phenotypes in *hbl-1(gf/*Δ*LCSs)* mutants. Our results indicate that the *C. elegans lin-28-lin-46* pathway regulates the temporal dynamics of nuclear accumulation of the HBL-1 transcription factor, conferring precision and robustness to the temporal down regulation of HBL-1 activity in parallel to *let-7*-family microRNAs.

## Results

### Deletion of genomic regions encompassing the *let-7* and *lin-4* complementary sites in the *hbl-1* 3’UTR results in extra seam cell phenotypes

To explore whether *lin-46* acts downstream of *lin-28* in the heterochronic pathway by modulating the regulation of *hbl-1* by *let-7* family and/or *lin-4* microRNA, we sought to generate *hbl-1(gf)* alleles free from regulation by these microRNAs. The *hbl-1* 3’UTR contains ten *let-7* complementary sites (LeCSs) and a single *lin-4* complementary site (LiCS; and collectively abbreviated as LCSs). In order to abrogate *let-7* and *lin-4* mediated regulation of *hbl-1*, we deleted a genomic region encompassing all LCSs in the *hbl-1* 3’UTR (*ma354;* Figure 1A, Table S2). We found that, similar to *mir-48/84/241(0)* mutants, *hbl-1(ma354[*Δ*LCSs])* animals have retarded seam cell defects, whereby L2 stage fates are reiterated at later stages, resulting in extra seam cells in young adult animals (Figure 1A).

Additionally, in our screens for large deletions in the progeny of CRISPR/Cas-9 injected animals, we recovered smaller 3’UTR deletions of various sizes that removed several but not all LeCSs in the *hbl-1* 3’UTR (Figure 1A and Table S2). We analyzed these smaller deletions along with the largest deletion (*ma354*), and found that most of these mutants also have extra seam cell phenotypes, although weaker than the *ma354* deletion (Figure 1A). 3’UTR deletions that removed more LeCSs resulted in stronger extra seam cell phenotypes, which is consistent with the idea that these LeCSs are functional and they act partially redundantly.

**Figure 1.**
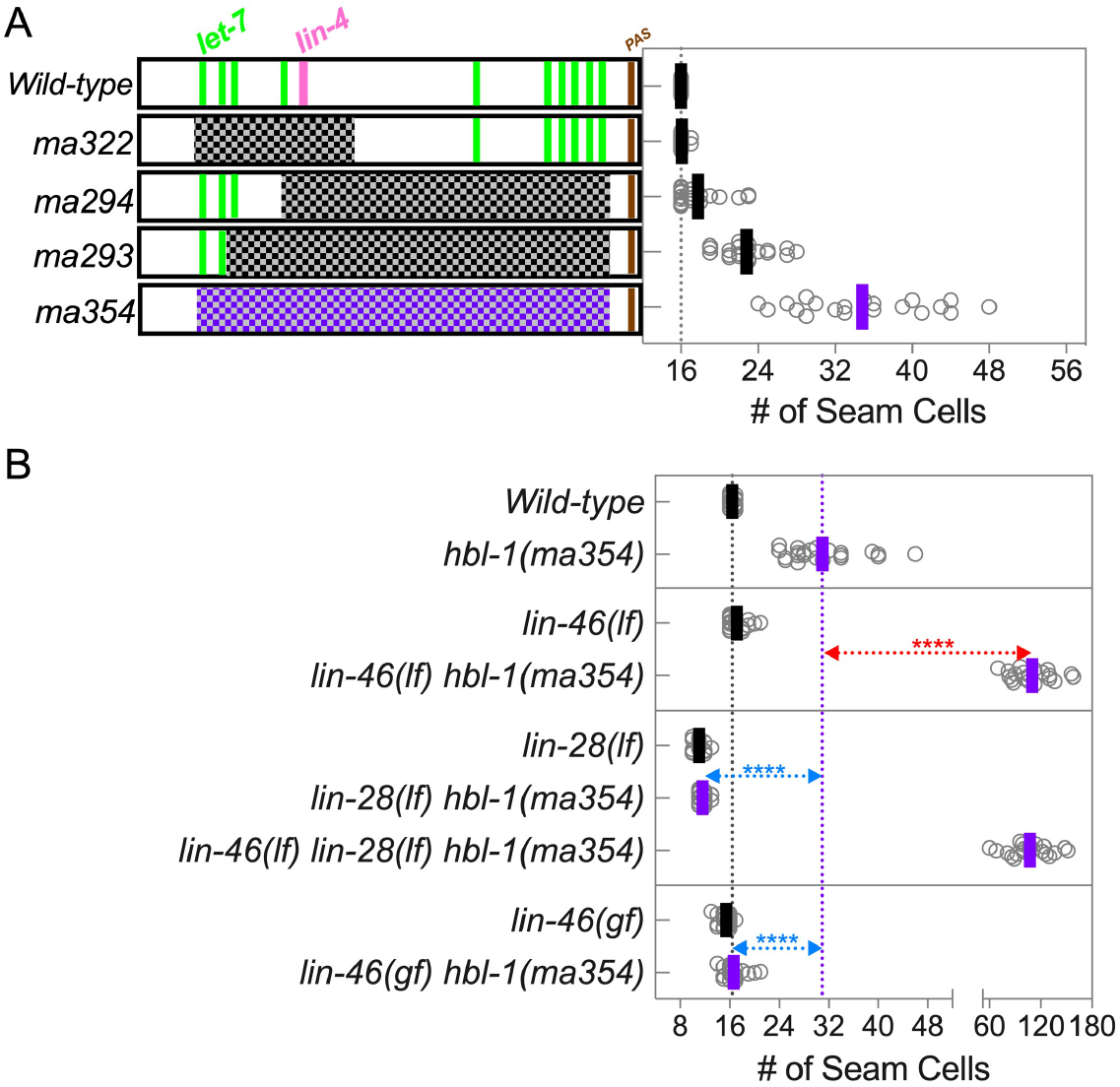
The *C. elegans lin-28-lin-46* pathway regulates L2 to L3 cellfate transitions independently of the *let-7* and *lin-4* complementary sites in the *hbl-1* 3’UTR. **A)** Deletion of genomic regions encompassing the *let-7* and *lin-4* complementary sites in the *hbl-1* 3’UTR results in extra seam cell phenotypes. The *hbl-1* 3’UTR contains ten *let-7* (green bars) and one *lin-4* (pink bar) complementary sites. Wildtype *hbl-1* 3’UTR and four deletion alleles are depicted on the y-axis and the number of seam cells observed in young adults of animals bearing these alleles are shown on the x-axis. Each dot shows the number of seam cells observed in a single animal and the lateral bars show the average seam number in the group of animals observed for each allele. **B)** Number of seam cells in single and compound mutants containing the *hbl-1(ma354)* allele are plotted. *lin-46* null allele enhances the extra seam cell phenotype of *hbl-1(ma354)* shown with red dotted line. *lin-28(lf)* suppresses the extra seam cell phenotype of *hbl-1(ma354)* and this suppression is *lin-46* dependent. *lin-46* gain-of-function allele similarly suppresses the extra seam cell phenotypes of *hbl-1(ma354)*. The Student’s t test is used to calculate statistical significance (p): n.s. (not significant) p > 0.05, *p < 0.05, **p < 0.01, ***p < 0.001, ****p < 0.0001.

### *lin-28(lf)* suppresses and *lin-46(lf)* enhances the extra seam cell phenotype of *hbl-1(ma354[*Δ*LCSs])*

Next, to test if the regulation of *hbl-1* by the *lin-28-lin-46* pathway was dependent on the *let-7* and *lin-4* complementary sites (LCSs) in the *hbl-1* 3’UTR, we generated compound mutants containing the *ma354* deletion, together with null alleles of *lin-28*, and/or *lin-46*. We found that *lin-28(lf)* suppresses, and *lin-46(lf)* substantially enhances, the extra seam cell phenotype of *hbl-1(ma354)* (Figure 1B), indicating that the regulation of *hbl-1* by *lin-28* does not require the LCSs in the *hbl-1* 3’UTR. We also found that *lin-46* is required for the suppression of *hbl-1(ma354)* by *lin-28(lf)* (Figure 1B). These results suggest that the *lin-28-lin-46* pathway regulates HBL-1 amount or activity through a mechanism independent of the *let-7/lin-4* regulation of HBL-1.

### A *lin-46(gf)* mutation can suppress the extra seam cell phenotypes of *hbl-1(ma354[*Δ*LCSs])*

LIN-46 is expressed precociously in *lin-28(lf)* animals [17] and so suppression of *hbl-1(gf)* by *lin-28(lf)* could be solely due to precocious LIN-46, which is sufficient to suppress L2 cell-fates and promote transition to L3 and later cell-fates [17]. To determine whether precocious LIN-46 expression alone is also sufficient to suppress the extra seam cell phenotypes of *hbl-1(ma354)* animals, we employed a *lin-46(gf)* mutation, *lin-46(ma467)*, that consists of a 12 bp deletion of *lin-46* 5’UTR sequences, and that results in precocious expression of LIN-46 and weak *lin-28(lf)*-like phenotypes [17]. We generated double mutant animals carrying *hbl-1(ma354)* and *lin-46(ma467)*, and found that the gain-of-function allele of *lin-46* suppresses the extra seam cell phenotypes of *hbl-1(ma354*) (Figure 1B), suggesting that precocious LIN-46 expression is sufficient to suppress the *hbl-1(gf)* phenotypes. This result supports the interpretation of the suppression of *hbl-1(gf)* by *lin-28(lf)* as resulting from precocious LIN-46 expression.

### Endogenously tagged HBL-1 is expressed in the nuclei of L1 and L2 stage hypodermal seam and hyp7 cells

The suppression and enhancement of the extra seam cell of *hbl-1(ma354)* by *lin-28(lf)* and *lin-46(lf)*, respectively, could reflect changes in the level of HBL-1 protein. In order to test for changes in the levels of HBL-1 protein in *lin-28(lf)* or *lin-46(lf)* mutants, we tagged *hbl-1* at the endogenous locus with mScarlet–I using CRISPR/Cas9 [12]. We observed that in wild type L1 and L2 larvae HBL-1::mScarlet–I localizes exclusively to the nucleus (Figure 2A, Table S1A), which is consistent with HBL-1 functioning as a transcription factor [18,19]. Moreover, consistent with previous reports, endogenously tagged HBL-1 is expressed in the wild type in the hypodermal seam and hyp7 cells of L1 and L2 stage larvae and it is not detected in L3 and L4 stage larvae [12,20] (Figure 2A, and Table S1A).

**Figure 2.**
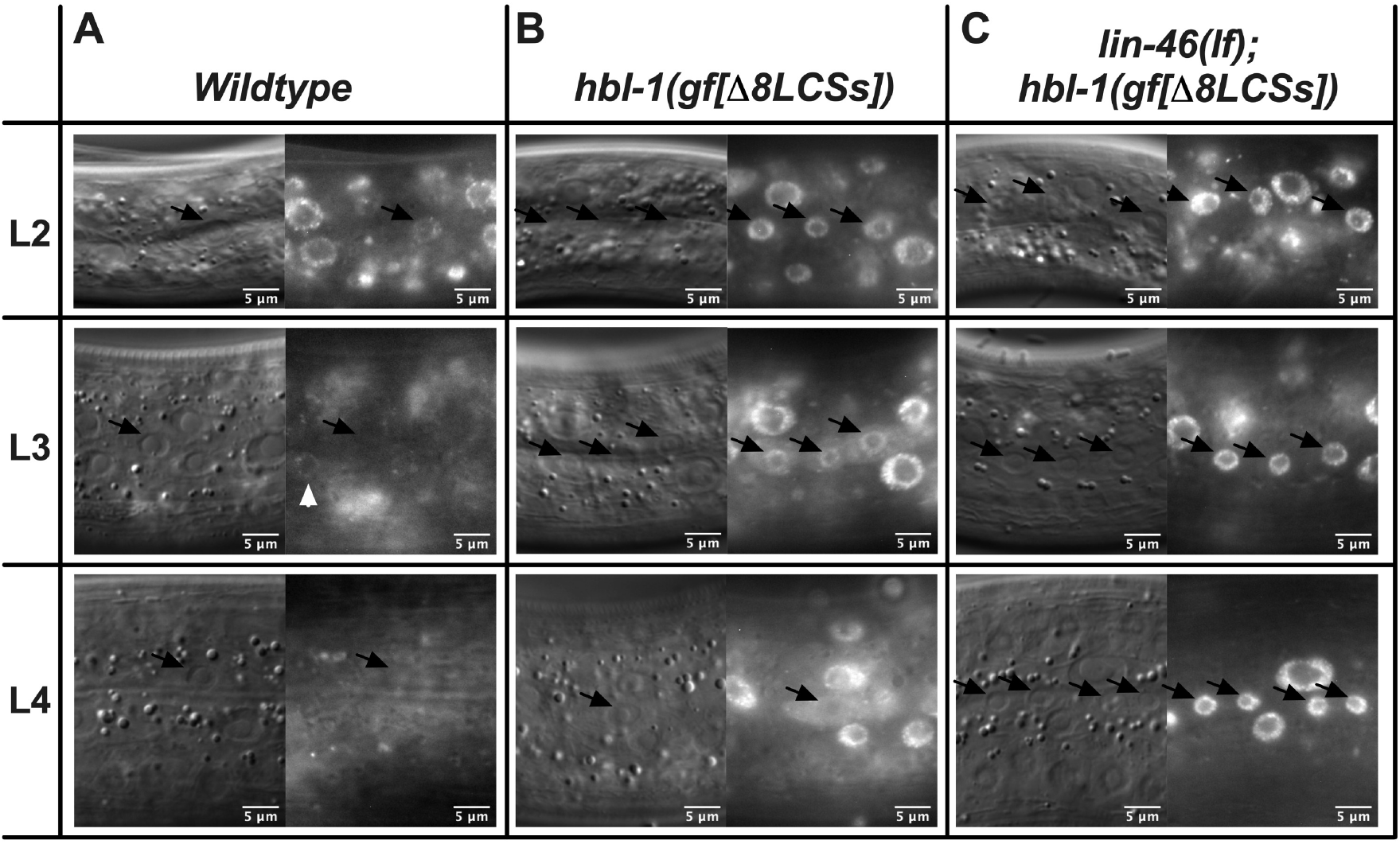
HBL-1 accumulates in the cytoplasm of L3 and L4 stage seam cells in larvae lacking LCSs in the *hbl-1* 3’UTR; and *lin-46* is required for this cytoplasmic accumulation of HBL-1. All seam cell nuclei are marked with black arrows. A) HBL-1 is expressed in hypodermal seam (black arrows) and hyp7 cells of L1 and L2 stage larvae and HBL-1 is not detected in L3 and L4 stage larvae. A rare occurrence of an L3 stage hyp7 cell expressing HBL-1 is marked with a white arrow head; and note that HBL-1 is absent in all other nuclei, including the seam nucleus (black arrow). B) In animals that lack a region of the *hbl-1* 3’UTR containing eight *let-7* complementary sites (LCSs), *hbl-1(ma430ma475)*, HBL-1 is present in hypodermal seam (black arrows) and hyp7 cells at all stages (L2-L4 shown). In these animals, HBL-1 accumulates in the cytoplasm of seam cells, at the L3 and L4 stages. L3 stage seam cells still display a marked nuclear HBL-1 accumulation whereas L4 stage seam cells display almost an equal distribution of HBL-1 in both the nucleus and the cytoplasm C). In animals lacking *lin-46* in addition to the eight LCSs in the *hbl-1* 3’UTR, HBL-1 does not accumulate in the cytoplasm of seam cells in L3 or L4 stage animals, rather HBL-1 accumulates in the nucleus of seam cells at all stages.

### HBL-1 is over-expressed and accumulates in the cytoplasm of L3 and L4 stage seam cells in larvae lacking LCSs in the *hbl-1* 3’UTR

To examine the impact of the loss of the LCSs in the *hbl-1* 3’UTR on the expression pattern of HBL-1, we deleted a region in the *hbl-1* 3’UTR of the mScarlet-I tagged *hbl-1* allele, generating the *ma430ma475 [hbl-1::mScarlet-I::*Δ*8LeCSs]* allele. The ma475 deletion encompasses eight LeCSs and the LiCS, similar to and only three base-pairs shorter than the *ma293* deletion (Figure 1A). Consistent with the previous reports that utilized GFP reporters fused with wildtype *hbl-1* 3’UTR [4,20], absence of these LeCSs results in HBL-1 expression that persists in the L3 and L4 stage hypodermal cells (Figure 2A&B). Interestingly, we observed that at the L3 and L4 stages, HBL-1 accumulates in the cytoplasm of the seam cells, which is accompanied by a reduction in the nuclear accumulation of HBL-1 (Figure 2B). This observation suggested that perhaps the nuclear accumulation of HBL-1 is hindered (or cytoplasmic accumulation of HBL-1 is facilitated) in these L3/L4 stage seam cells.

### *lin-28* is required for the nuclear accumulation of HBL-1 in the seam cells of L2 stage larvae

*lin-28(lf)* animals skip L2 stage proliferative seam cell-fates, suggesting that *lin-28* is required to support the activity of HBL-1 in the L2. Moreover, loss of *lin-28* can suppress the extra seam cell phenotypes of *hbl-1* gain-of-function (gf) mutants (Figure 1B), indicating that *lin-28* is also required to support high and/or prolonged expression of HBL-1. We observed that in double mutants containing *hbl-1(ma430ma475)* and *lin-28(lf)*, HBL-1 accumulates primarily in the cytoplasm and is largely absent from the nucleus of the L2 stage seam cells (Figure 3B), which explains the lack of HBL-1 activity and the suppression of *hbl-1(gf)* extra seam cell phenotypes in larvae lacking *lin-28*. In order to test the possibility of an effect of the linker or the mScarlet-I tag on the localization of HBL-1, we also tagged another transcription factor (whose nuclear accumulation is not affected by *lin-28(lf);* Figure S1A), *daf-12*, at its endogenous locus with the same linker and mScarlet-I and asked if the localization of DAF-12 changed in *lin-28(lf)* animals (Figure S1). We observed that, unlike HBL-1, linker-mScarlet tagged DAF-12 did not accumulate in the cytoplasm of L2 stage seam cells in *lin-28(lf)* animals (Figure S1B), suggesting that the linker-mScarlet-I tag could not be the cause of cytoplasmic accumulation of HBL-1.

**Figure 3.**
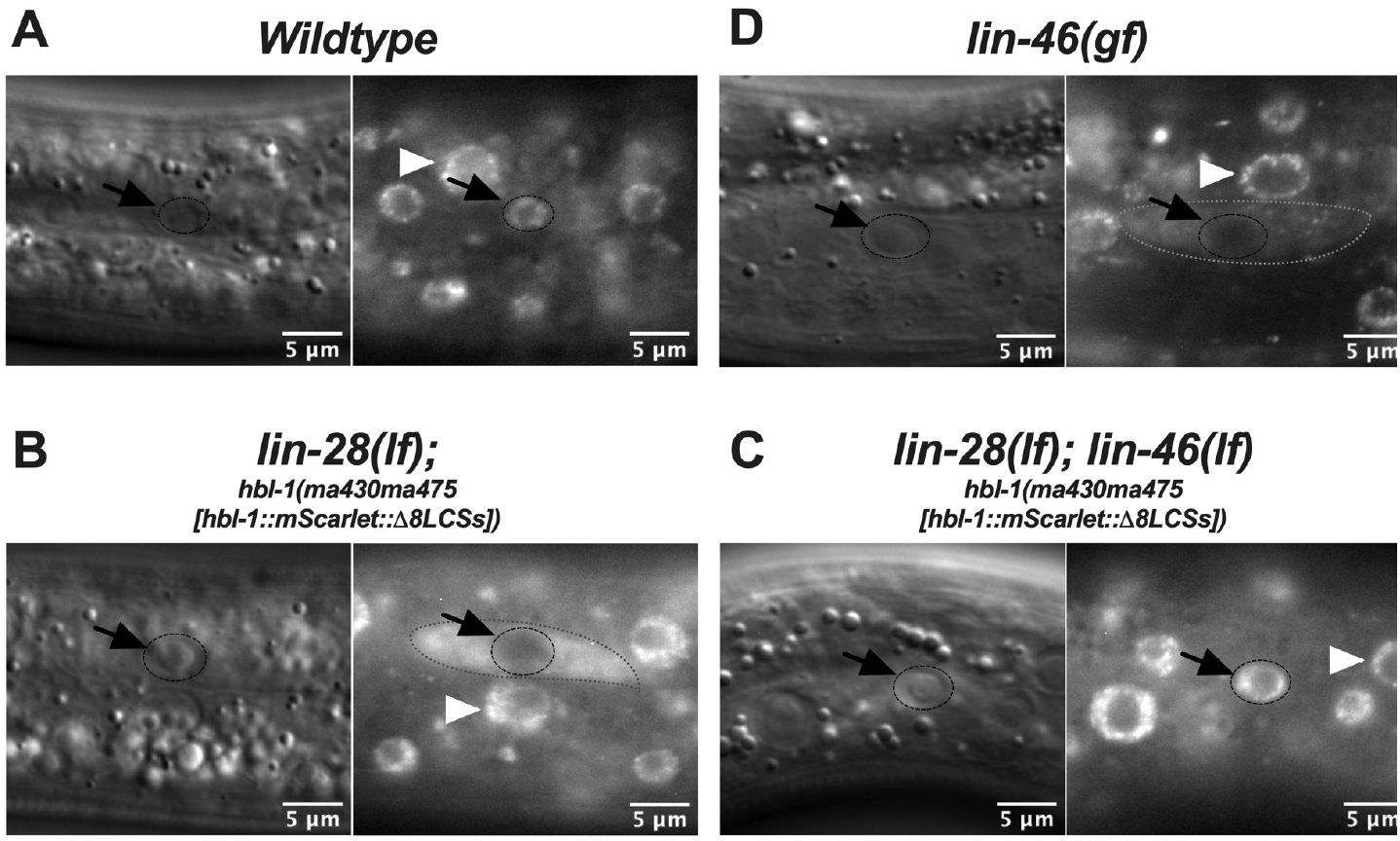
The *lin-28-lin-46* pathway regulates the nuclear accumulation of HBL-1. DIC and fluorescent images of hypodermal (seam and hyp7) cells in L2 stage larvae. Black arrows and dashed circles show seam nuclei and white arrow heads show hyp7 nuclei. A) HBL-1 accumulates in the nucleus in wildtype animals. B) *lin-28* is required for the nuclear accumulation of HBL-1 in the seam cells of L2 stage larvae. HBL-1 is dispersed in the cytoplasm (outlined by dashed line) of L2 stage seam cells in *lin-28(lf)* animals. Note that HBL-1 still accumulates in hyp7 nuclei (e.g. whit arrowhead). C) *The lin-28* target *lin-46* is required to prevent the nuclear accumulation of HBL-1 in L2 stage seam cells of *lin-28(lf)* larvae. D) Precocious/ectopic LIN-46 expression is sufficient to reduce the nuclear accumulation of HBL-1 in the seam cell of L2 stage larvae. HBL-1 is present in the cytoplasm (outlined by dashed line) of the seam cell in the picture. Note that HBL-1 still accumulates in hyp7 nuclei (e.g. whit arrowhead).

### *lin-46* activity is required for the cytoplasmic accumulation of HBL-1 both in *hbl-1(ma430ma475)* and *lin-28(lf)* animals

HBL-1 accumulates both in the nucleus and the cytoplasm of L3 and L4 stage seam cells in *hbl-1(ma430ma475[mScarlet-I::*Δ*8LCSs])* larvae (Figure 2B), when we combined *hbl-1(ma430ma475[mScarlet-I::*Δ*8LCSs]) with lin-46(lf)*, we did not observe cytoplasmic accumulation of HBL-1 at the L3 and L4 stages, rather HBL-1 accumulates in the nucleus at all stages (Figure 2C), indicating that the L3/L4 stage cytoplasmic accumulation of HBL-1 requires *lin-46* activity.

HBL-1 accumulates primarily in the cytoplasm of L2 stage seam cells in *lin-28(lf)* larvae (Figure 3B). By contrast, in L2 larvae lacking both *lin-28* and *lin-46*, HBL-1 no longer accumulates in the cytoplasm of L2 or later stage seam cells, rather HBL-1 accumulates in the nucleus of the seam cells at all stages (Figure 3C and Table S1).

In brief, *lin-46(lf)* results in the loss of cytoplasmic HBL-1 accumulation at all stages, accompanied by restoration of nuclear accumulation of HBL-1 (Table S1).

### Precocious LIN-46 expression if sufficient to reduce the nuclear accumulation of HBL-1

Lastly, we found that the precocious LIN-46 expression, which is sufficient to suppress the extra seam cell phenotypes of *hbl-1(ma354)* (Figure 1B) is also sufficient to reduce the nuclear accumulation of HBL-1 (Figure 3D).

These results, together with those presented above show that the suppression of *hbl-1(ma354)* phenotypes by *lin-28(lf)* or *lin-46(gf)* is accompanied by increased cytoplasmic accumulation of HBL-1 and reduction in the nuclear accumulation of HBL-1. These findings suggest that LIN-46 negatively regulates HBL-1 activity by hindering its nuclear accumulation.

### Spatiotemporal occurrence of cytoplasmic HBL-1 accumulation coincides with LIN-46 expression

Endogenously tagged *lin-46* is expressed in the seam cells of L3&L4 stage larvae but not in the seam cells of L1&L2 stage larvae [17]. Therefore, the onset of LIN-46 expression coincides with the onset of cytoplasmic accumulation of HBL-1 in *hbl-1(ma430ma475[mScarlet-I::*Δ*8LCSs])* animals (which are, unlike the wildtype, not capable of properly down regulating HBL-1 at the end of the L2 stage owing to the lack of LCSs). In *lin-28(lf)* animals, LIN-46 is expressed precociously [17], starting in mid L1 stage seam cells. This precocious onset of LIN-46 expression in *lin-28(lf)* animals also coincides with the precocious onset of the cytoplasmic accumulation of HBL-1 in *lin-28(lf)* animals.

We note that LIN-46 expression is detected in the seam cells but not in the hyp7 cells [17]; and the presence/absence of LIN-46 expression in these two hypodermal cell types correlates with the presence/absence of cytoplasmic accumulation of HBL-1. For example, whereas HBL-1 accumulates in both the cytoplasm and the nucleus of seam cells L3/L4 stage *hbl-1(ma430ma475[mScarlet-I::*Δ*8LCSs])* animals, HBL-1 accumulates only in the nucleus of the hyp7 cells (Figure 2B). Similarly, whereas HBL-1 accumulates primarily in the cytoplasm of L2 stage seam cells of *lin-28(lf)* animals, HBL-1 still accumulates in the nucleus of hyp7 cells in these animals (Figure 3B, black vs white arrows).

In brief, cytoplasmic accumulation of HBL-1 is observed in the hypodermal cells where (seam) and when [by the L3 stage in wild-type and by the late L1 stage in *lin-28(lf)*] LIN-46 is expressed (Table S1).

## Discussion

Our results suggest that the *C. elegans lin-28-lin-46* pathway regulates the nuclear accumulation of HBL-1, a transcription factor that specifies L2 stage proliferative cell-fates and opposes the progression to L3 stage self-renewal cell-fates during *C. elegans* development. *lin-28* is required for the nuclear accumulation of HBL-1 in the hypodermal seam cells at the L2 stage, which is, in turn, necessary for the execution of L2 stage proliferative cell-fates. The *lin-28* target *lin-46* is responsible for preventing the nuclear accumulation of HBL-1 in *lin-28(lf)* animals; and in wildtype animals *lin-28*-mediated repression of the LIN-46 expression at the L1 and L2 stages [17] allows the nuclear accumulation of HBL-1 at those early larval stages. Using a *lin-46* 5’ UTR mutation that render LIN-46 expression poorly repressed by *lin-28*, we show that precocious LIN-46 expression in the seam cells of the L1/L2 stage larvae is sufficient to reduce the nuclear accumulation of HBL-1. Furthermore, using *hbl-1* gain-of-function mutations with *let-7* and *lin-4* sites deleted from the *hbl-1* 3’ UTR, we show that the *lin-28-lin-46* pathway acts in parallel to *let-7* family microRNAs. Hence, these two parallel pathways – the microRNA pathway controlling the rate of synthesis of HBL-1 through repression of *hbl-1* mRNA translation, and the *lin-28-lin-46* pathway controlling the nuclear accumulation of HBL-1 – function together to ensure the precision and robustness of stage-specific HBL-1 down regulation (Figure 4A).

**Figure 4.**
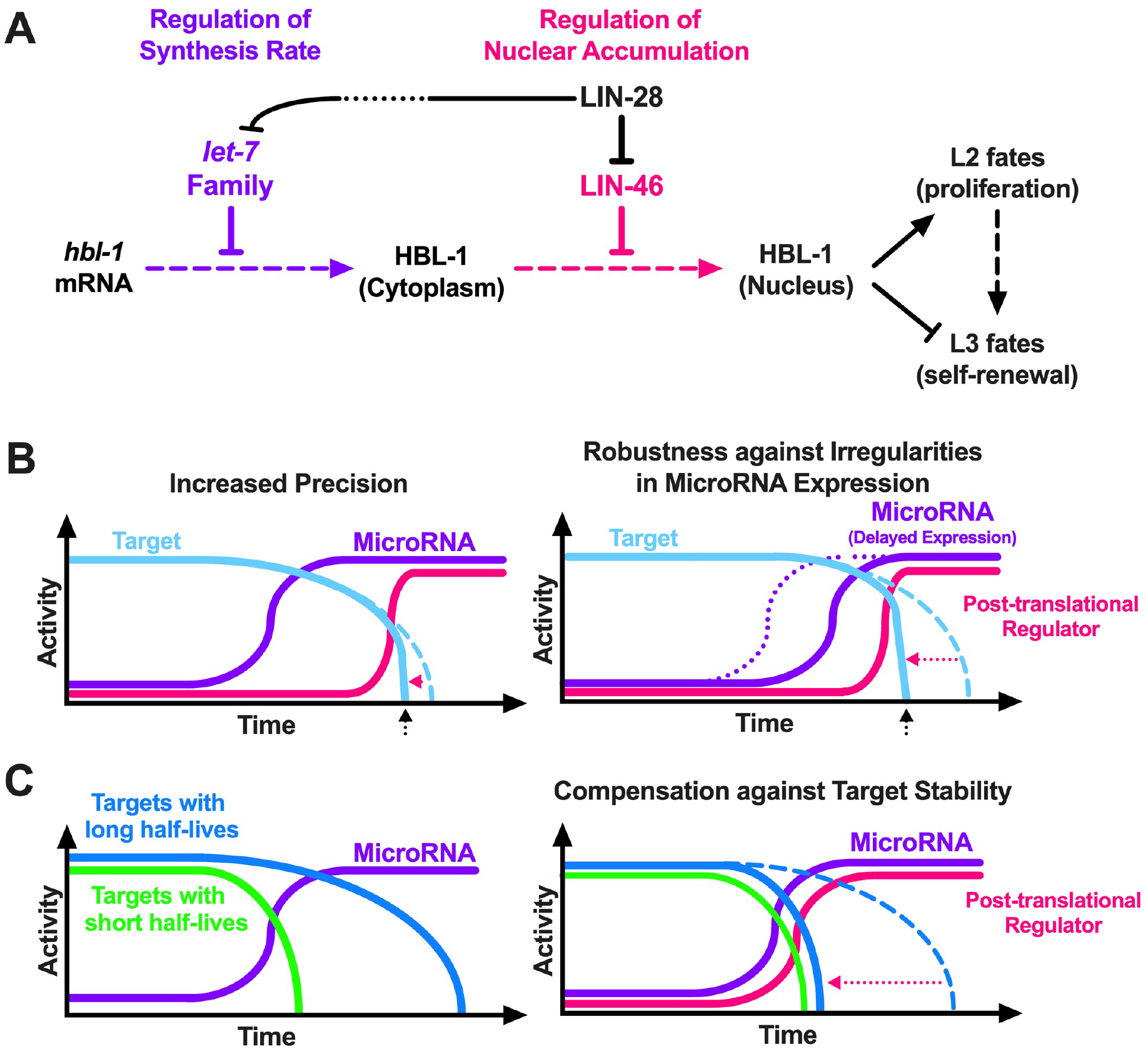
Regulation of gene activity through microRNA-mediated repression of translation accompanied by post-translational regulation of microRNA targets. **A)** The conserved RNA binding protein LIN-28 indirectly (indicated by the dotted line break) regulates the transcription [33] and activities [34] of *let-7*-family (*mir-48, mir-84, mir-241*) microRNAs, which inhibit the synthesis of HBL-1. LIN-28 also represses the expression of LIN-46 [17], which controls the nuclear accumulation of HBL-1. Temporal down-regulation of LIN-28 at the end of the L2 stage allows LIN-46 to accumulate, which acts together with the *let-7*-family microRNAs to ensure precise and robust temporal down-regulation of HBL-1 activity. **B)** Hypothetical activity trajectories of a microRNA and its target(s) against time are plotted. Dashed lines represent the trajectory in the absence of the hypothetical post-translational regulator. A post-translational regulator of a microRNA target can increase the precision of temporal down-regulation of this target (B, left) or confer robustness against irregularities in microRNA expression (B, right). This second scenario is similar to what is thought to happen in *C. elegans* larvae developing in the presence of pheromones or other L2d-inducing conditions: *let-7*-family expression is delayed [14,15] and LIN-46 activity becomes more important for down-regulating HBL-1 [12]. Thus, LIN-46 confers robustness against physiological delays in the expression of *let-7*-family microRNAs. **C)** Post-translational regulators of microRNA targets can compensate against target stability. In this scenario, the activities of targets with short half-lives (green) are sharply reduced as the targeting microRNA activity increases, whereas the activities of targets with long half-lives (blue) are maintained even during a period of time after the targeting microRNA is fully activated (C, Left). A dual activation of a microRNA and a post-translational regulator of its targets with long half-lives would enable a faster reduction in the activities of these targets, making sharp transitions from active to inactive states possible for these more stable microRNA targets (C, right).

In wildtype animals HBL-1 and LIN-46 are expressed at temporally distinct stages: HBL-1 is expressed at the L1&L2 stages whereas LIN-46 is expressed at the L3&L4 stages. In larvae of certain mutants, such as *hbl-1(gf)* L3 and L4 larvae, or *lin-28(0)* L1 and L2 larvae, LIN-46 and HBL-1 expression overlap, and cytoplasmic accumulation of HBL-1 is observed, accompanied by a reduction in the nuclear accumulation of HBL-1. Our data further show that the nucleus-to-cytoplasm displacement of HBL-1 in these contexts depends on *lin-46* activity. Therefore, one might have expected to observe cytoplasmic accumulation of HBL-1 after the L2-to-L3 transition in wild type larvae, when LIN-46 begins to accumulate. Curiously, in wild-type animals, cytoplasmic accumulation of HBL-1 is not evident at any stage, despite the presence of LIN-46 in L3-adult animals. If LIN-46 causes cytoplasmic accumulation of HBL-1 in the wild type, why is HBL-1 not detected in L3 and L4 larvae? One explanation could be that in wildtype larvae, the post-translational repression of HBL-1 activity by LIN-46 (via cytoplasmic localization) functions semi-redundantly with the translational repression of HBL-1 by *let-7* family microRNAs. In this scenario, the microRNA pathway could exert the lion’s share of HBL-1 down regulation at the L2-to-L3 transition, and the up regulation of LIN-46 at the L3 could play a secondary role to inhibit the nuclear accumulation of any residual HBL-1 protein. Indeed, in support of this idea, a low level of HBL-1 expression persists in the nuclei of L3 stage seam cells in *lin-46(lf)* animals (Table S1). This scenario is also consistent with the differing strengths of the weaker retarded phenotypes of *lin-46(lf)* animals compared to *hbl-1(gf)* animals under standard culture conditions (Figure 1B).

Interestingly, conditions such as diapause-inducing stress signals that enhance *lin-46(lf)* phenotypes [12] also result in a reduction in the expression of *let-7* family microRNAs [15], resulting in a shift from primarily-microRNA mediated regulation of HBL-1 to primarily LIN-46 mediated regulation [12]. In this context, where wildtype larvae experience diapause-inducing stress signals, and hence LIN-46-mediated cytoplasmic localization becomes the primary mode of HBL-1 down regulation, we expected to observe cytoplasmic accumulation of HBL-1 after the L2-to-L3 transition. However, we could not detect any cytoplasmic accumulation of HBL-1 in wildtype animals under such conditions. There are several possible explanations for this, including the possibility that the intrinsically low HBL-1 abundance in the wildtype renders dispersed cytoplasmic HBL-1 to be simply below the limit of detection in our fluorescence microscopy assays.

Sequence homology places LIN-46 into a conserved protein family whose members include bacterial MOEA as well as human GPHN (Gephyrin), which are implicated in molybdenum cofactor (MoCo) biosynthesis [21]. GPHN is also reported to function as a scaffold protein that is required for clustering of neurotransmitter receptors [22,23], and is shown to physically interact with several other proteins [24], including tubulin [25], dynein [26], and mTOR [27]. It is not known if LIN-46 possesses MOEA-related enzymatic activity and/or has scaffolding functions similar to GPHN, and if so, how such activities could (directly or indirectly) contribute to inhibiting the nuclear accumulation of a transcription factor such as HBL-1.

Analysis of HBL-1 amino acid sequence does not reveal a predicted nuclear localization signal (NLS) that could mediate HBL-1 nuclear transport. If HBL-1 has an unconventional or “weak” NLS, it is possible that other unknown factors may be required to efficiently couple HBL-1 to the nuclear import machinery. LIN-46 might inhibit HBL-1 nuclear accumulation by binding or modifying a factor critical for HBL-1 nuclear transport. Alternatively, LIN-46 could bind or modify HBL-1 directly so as to prevent its association with the nuclear import machinery. It is also possible that LIN-46 could act not by preventing nuclear import of HBL-1, but by causing HBL-1 to be trapped in the cytoplasm, for example through the formation of LIN-46-HBL-1 complexes in association with a cytoplasmic compartment.

Regulation of nuclear accumulation in the context of temporal cell-fate specification during *C. elegans* development has not previously been reported. Other transcription factors, including LIN-14, DAF-12 [28], and LIN-29, play key roles in regulating temporal cell-fates during *C. elegans* development. These other transcription factors are also regulated by microRNAs, like HBL-1, and regulation of their temporal abundances is important for the proper execution of stage-specific cell-fates. Although we have no evidence that LIN-46 may also regulate the nuclear/cytoplasmic partitioning of other heterochronic pathway proteins, our findings suggest that similar post-translational mechanisms might be in place to function in parallel with the microRNA-mediated regulation and hence promote the robust temporal regulation of key developmental regulators.

Many *let-7* targets, as well as many targets of other microRNAs, in worms, flies, and mammals are transcription factors [29–32]. Therefore, similar mechanisms, whereby a transcription factor is regulated both by a microRNA and in parallel by a gene product that controls the nuclear accumulation of the same transcription factor, may be common. Additionally, the regulation of the *let-7* microRNA by LIN-28 is widely conserved. Targets of LIN-28 in other species may have roles, similar to LIN-46, to regulate *let-7* targets, controlling their nuclear accumulation in particular and their activities by means of post-translational interventions in general. A dual control of a gene product – its synthesis rate by microRNAs and activity by post-translational regulators – would allow more precise and/or more robust transitions between active to inactive states (Figure 4B), or sharper transitions when proteins that have long half-lives are involved (Figure 4C).

## Materials and Methods

### *C. elegans* culture conditions

*C. elegans* strains used in this study and corresponding figures in the paper are listed in Table S3. *C. elegans* strains were maintained at 20°C on nematode growth media (NGM) and fed with the *E. coli* HB101 strain.

### Assaying extra seam cell phenotypes

The worms were scored at the young adult stage (determined by the gonad development) for the number of seam cells using fluorescence microscopy with the help of the *maIs105* [*pCol-19::gfp*] transgene that marks the lateral hypodermal cell nuclei or the *wIs51[pScm::gfp*] transgene that marks the seam cell nuclei.

Each circle on the genotype versus number of seam cells plots shows the observed number of seam cells on one side of a single young adult worm. More than 20 worms for each genotype or condition are analyzed and the average number of seam cells are denoted by lateral bars in the genotype versus number of seam cell plots. The Student’s t test is used to calculate statistical significance when comparing different genotypes or conditions. The GraphPad Prism 8 software is used to plot the graphs and for statistical analysis.

### Microscopy

All DIC and fluorescent images are obtained using a ZEISS Imager Z1 equipped with ZEISS Axiocam 503 mono camera, and the ZEN Blue software. Prior to imaging, worms were anesthetized with 0.2 mM levamisole in M9 buffer and mounted on 2% agarose pads. The ImageJ Fiji software is used to adjust the brightness and contrast of the images to enhance the visualization of the fluorescent signal. All images are taken using the same microscopy settings and a standard exposure time for all larval stages and genetic background, but because the brightness and contracts of the individual images are enhanced separately, the signal intensities do not represent the relative expression levels and cannot be used to compare expression levels across different larval stages of genetic backgrounds.

### Generation of new alleles using CRISPR/Cas9

CRISPR/Cas9 genome editing tool are used to generate the *hbl-1* 3’UTR deletion alleles, the *lin-46* open-reading frame (ORF) deletion allele, and to tag the *daf-12* gene with GFP and mScarlet-I at its endogenous locus.

For the *hbl-1* 3’UTR deletions and the *lin-46* ORF deletion (Table S2), a mixture of plasmids encoding SpCas9 (pOI90), and a pair of single guide RNAs (sgRNAs, expressed from pOI83 [12]) targeting both sites of interest (primers: Table S4) and the *unc-22* gene (pOI91) as co-CRISPR marker [35], and a rol-6(su1006) containing plasmid (pOI124) as co-injection marker was injected into the germlines of young adult worms. F1 roller and/or twitcher animals (50+ worms until the desired allele is detected) were cloned and screened by PCR amplification (primers: Table S4) for the presence of the expected size PCR product consistent with deletion of the genomic region spanning between the sites targeted by the pair of two guides.

To tag *daf-12* at the endogenous locus with the same linker and mScarlet-I sequence as the *hbl-1*(ma430) allele, a homologous recombination (HR) donor plasmid (pOI193) and sgRNA plasmid (pOI93) were included in the CRISPR mix that contained plasmids pOI90 (spCas9), pOI91 (unc-22 guide), pOI124 (rol-6). The HR plasmid pOI193 contains C-terminal end of *<u>daf-12</u>* fused in frame with the linker and mScarlet-I sequence that was subcloned from pOI191, which was used to tag *hbl-1* to generate the *ma430* allele. To tag *daf-12* with GFP instead of pOI193, an HR donor plasmid (pOI122) that contained GFP sequence flanked by HR sequences was included in the CRISPR mix.

In all new CRISPR alleles, genomic regions spanning the deletion site or the HR arms and the tags introduced were sequenced using Sanger sequencing. For each allele, a single worm with a precise (HR) edited locus was cloned and backcrossed twice before used in the experiments.

## Supplemental Figures

**Figure S1.**
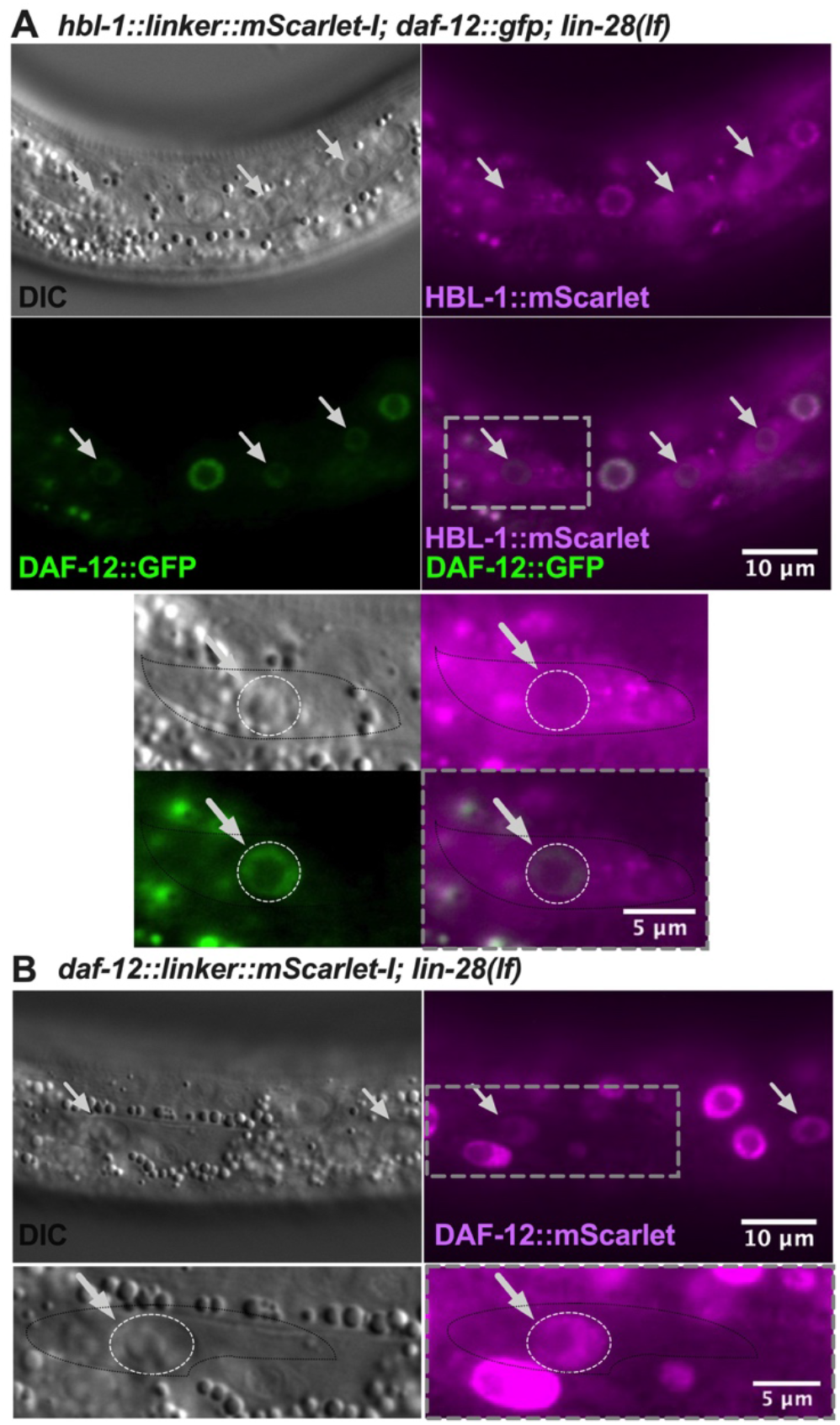
Nuclear localization of GFP or mScarlet-I tagged DAF-12 is not regulated by *lin-28*. DIC and fluorescent images showing hypodermal seam and hyp7 nuclei of L2 stage *lin-28(lf)* larvae. White arrows indicate the seam cell nuclei. A) In the absence of *lin-28(lf)*, HBL-1 is majorly excluded from the nuclei of L2 stage seam cells whereas nuclear localization of DAF-12 (GFP tagged) is not affected. B) The linker and mScarlet-I tag that was used to tag *hbl-1* do not have an effect on nuclear localization of DAF-12 in the *lin-28(lf)* background.

## Supplemental Tables

**Table S1.**
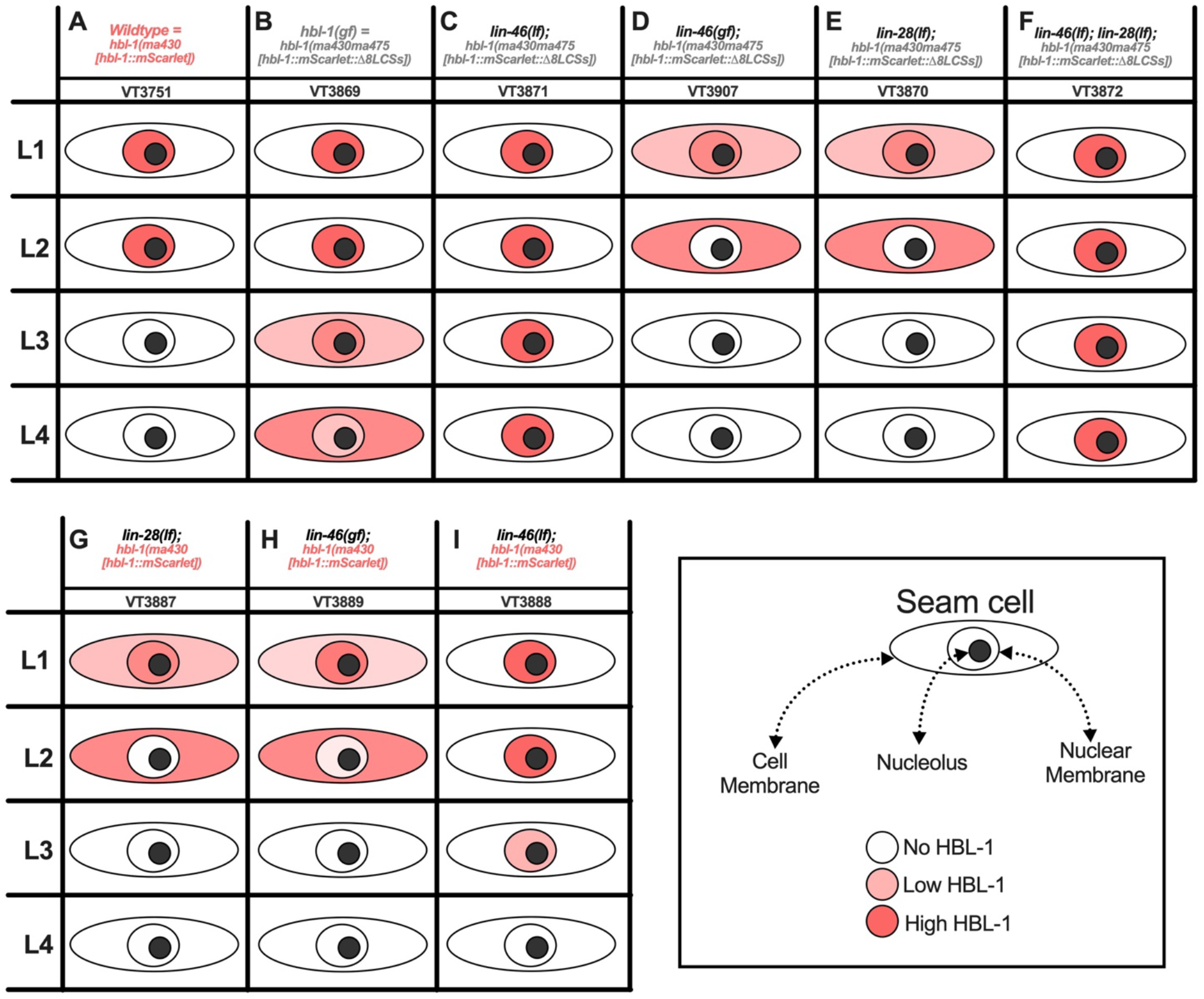
Schematic representations of nucleo-cytoplasmic localization of HBL-1 in hypodermal seam cells across four larval stages and various genetic backgrounds. HBL-1 expression (pink color) is denoted in nuclear/cytoplasmic compartments of seam cells for each larval stage and genotype. Ten animals for each larval stage and genotyped were analyzed. Genotypes and corresponding strain names (VT numbers) are denoted at the top of each column.

**Table S2.**
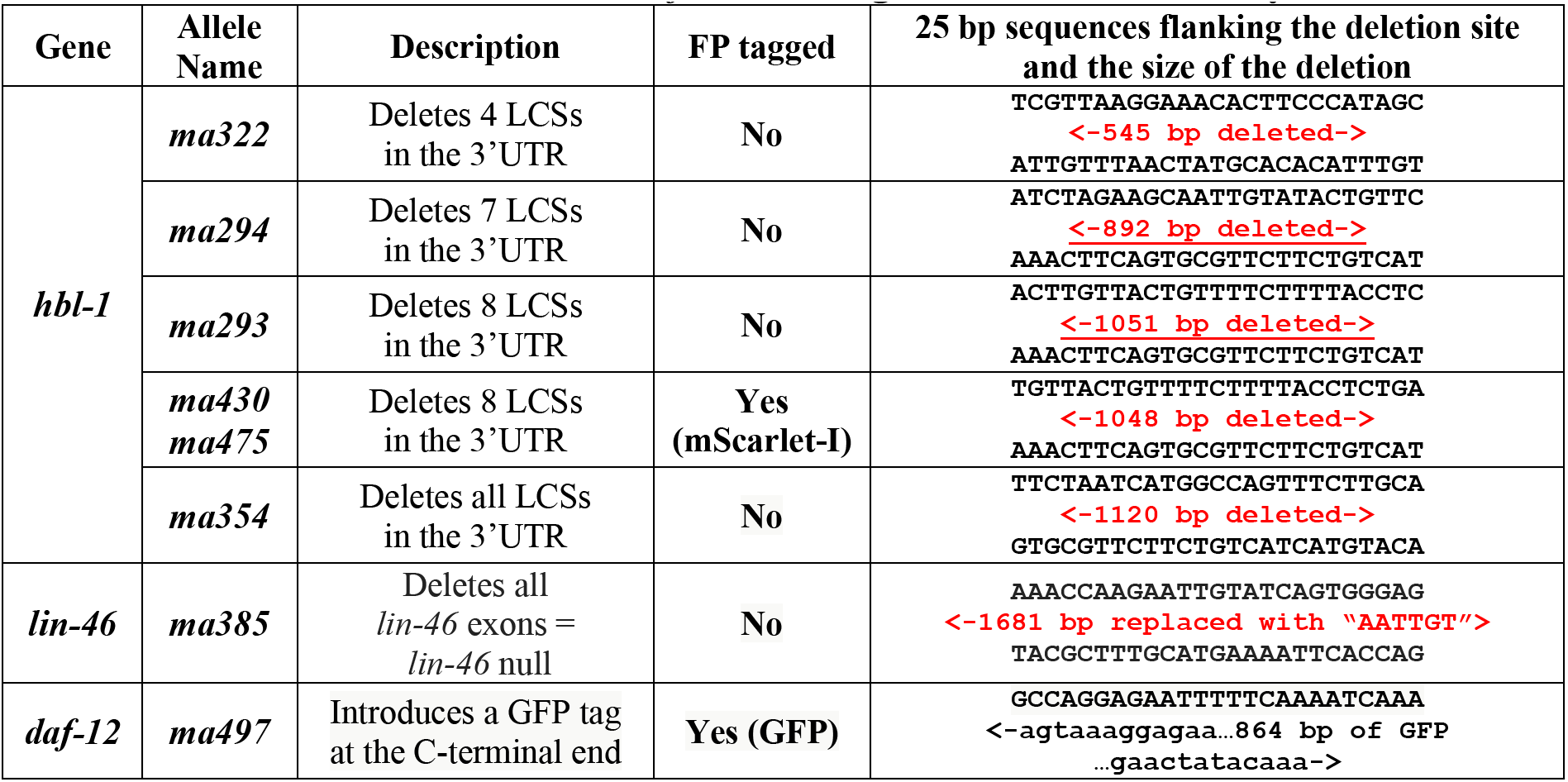
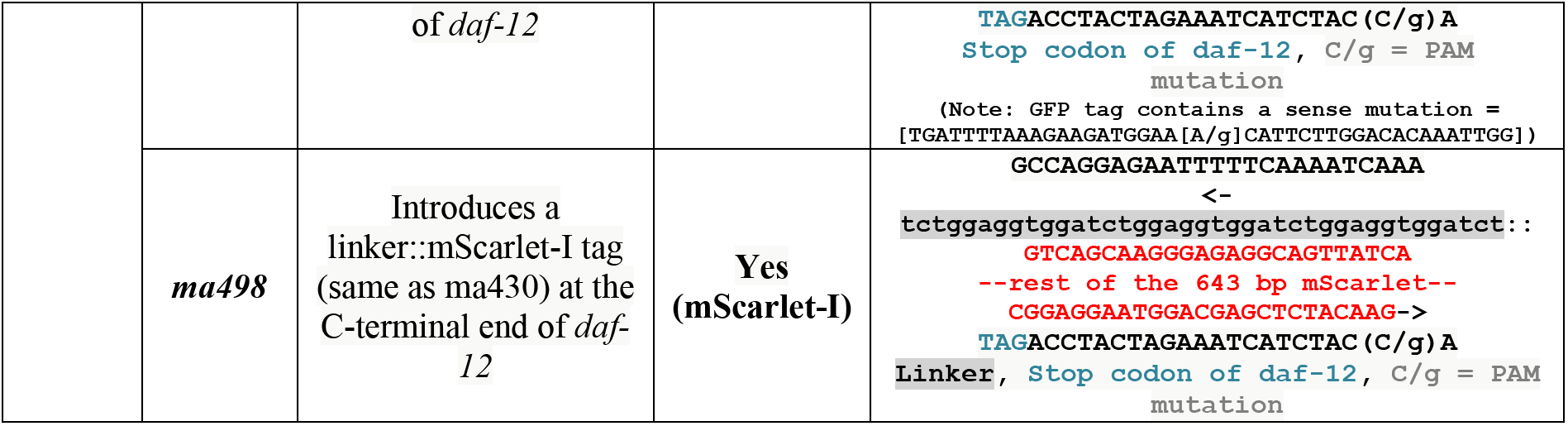
List of new *hbl-1, lin-46, daf-12* alleles generated for this study

**Table S3.** *C. elegans* strains used in this study.

| Strain name | Genotype | Related figures |
| --- | --- | --- |
| VT1367 | *maIs105 V* | Figure 1A&B |
| VT3307 | *maIs105 V; hbl-1(ma294) X* | Figure 1A |
| VT3336 | *maIs105 V; hbl-1(ma293) X* | Figure 1A |
| VT3399 | *maIs105 V; hbl-1(ma332) X* | Figure 1A |
| VT3500 | *wIs51 V; hbl-1(ma354) X* | Figure 1A&B |
| VT3593 | *lin-46(ma385) I; maIs105 V* | Figure 1B |
| VT3696 | *lin-46(ma385) wIs51 V; hbl-1(ma354) X* | Figure 1B |
| VT790 | *lin-28(n719) I; maIs105 V* | Figure 1B |
| VT3571 | *lin-28(n719) I; maIs105 V; hbl-1(ma354) X* | Figure 1B |
| VT3696 | *lin-46(ma385) wIs51 V; hbl-1(ma354) X* | Figure 1B |
| VT3698 | *lin-28(n719) I; lin-46(ma385) wIs51 V; hbl-1(ma354) X* | Figure 1B |
| VT3855 | *lin-46(ma467) maIs105 V* | Figure 1B |
| VT3891 | *lin-46(ma467) maIs105 V; hbl-1(ma354) X* | Figure 1B |
| VT3751 | *maIs105 V; hbl-1(ma430[hbl-1::mScarlet-I]) X* | Figure 2A&3A, Table S1A |
| VT3870 | *lin-28(n719) I; maIs105 V; hbl-1(ma430ma475) X* | Figure 2B, Table S1E |
| VT3872 | *lin-28(n719) I; wIs51 lin-46(ma385) V; hbl-1(ma430ma475) X* | Figure 2C, Table S1F |
| VT3889 | *lin-46(ma467) maIs105 V; hbl-1(ma430) X* | Figure 2D, Table S1H |
| VT3869 | *wIs51 V; hbl-1(ma430ma475[hbl-1::mScarlet-I::UTRdel]) X* | Figure 3B, Table S1B |
| VT3871 | *wIs51 lin-46(ma385) V; hbl-1(ma430ma475) X* | Figure 3C, Table S1C |
| VT3907 | *lin-46(ma467) maIs105 V; hbl-1(ma430ma475) X* | Table S1D |
| VT3887 | *lin-28(n719) I; maIs105 V; hbl-1(ma430) X; syls(ajm-1::gfp)* | Table S1G |
| VT3888 | *lin-46(ma385) maIs105 V; hbl-1(ma430) X; syls(ajm-1::gfp)* | Table S1I |
| VT3922 | *lin-28(n719) I; daf-12(ma497[daf-12::gfp] hbl-1(ma430[hbl-1::mScarlet-I]) X* | Figure S1 |
| VT3924 | *lin-28(n719) I; daf-12(ma498[daf-12::mScarlet-I]) X* | Figure S1 |

**Table S4.**
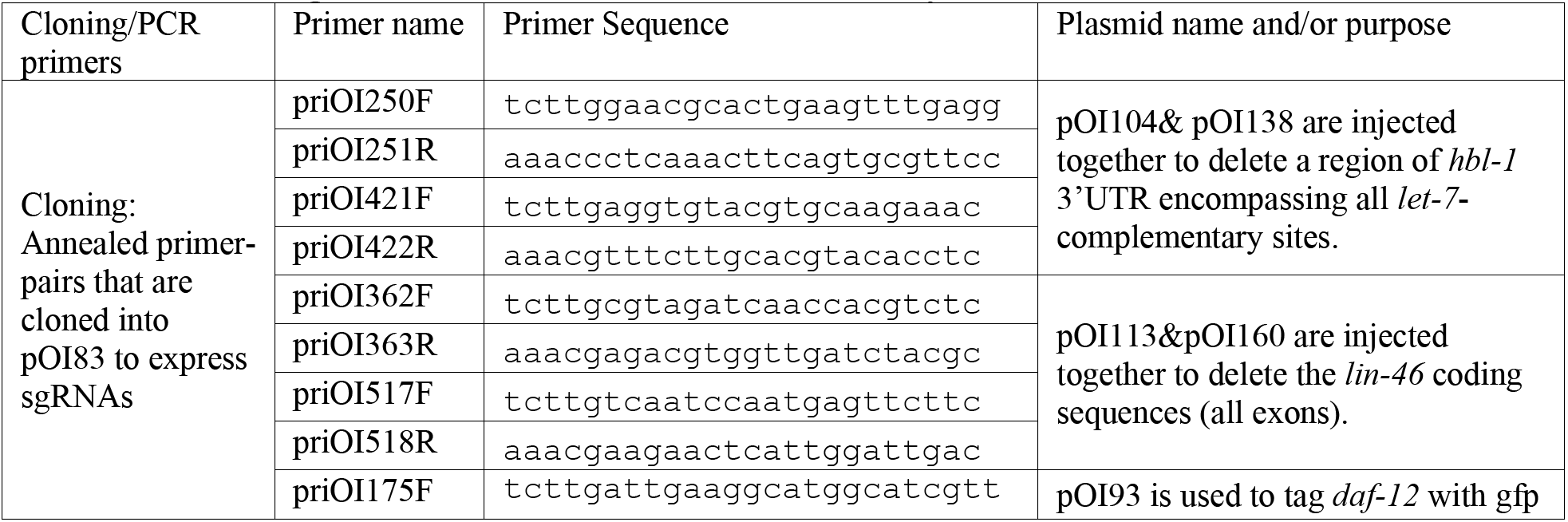
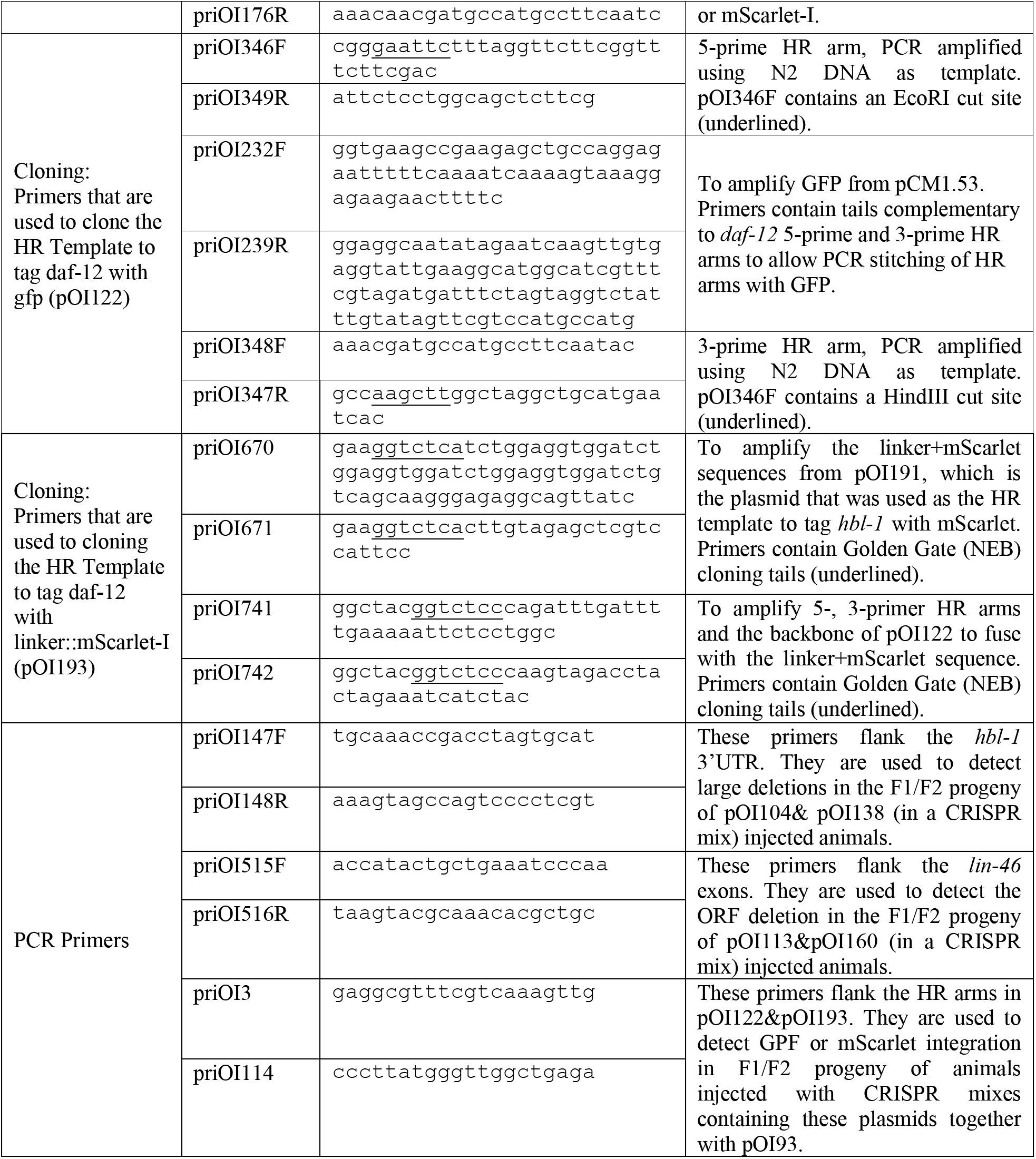
Cloning and PCR Primers used in this study.

## REFERENCES

1. Sulston, J.E., and Horvitz, H.R. (1977). Post-embryonic cell lineages of the nematode, Caenorhabditis elegans. Dev. Biol. 56, 110–156.

2. Ambros, V., and Horvitz, H.R. (1984). Heterochronic mutants of the nematode Caenorhabditis elegans. Science 226, 409–416.

3. Lee, R.C., Feinbaum, R.L., and Ambros, V. (1993). The C. elegans heterochronic gene lin-4 encodes small RNAs with antisense complementarity to lin-14. Cell 75, 843–854.

4. Abbott, A.L., Alvarez-Saavedra, E., Miska, E.A., Lau, N.C., Bartel, D.P., Horvitz, H.R., and Ambros, V. (2005). The let-7 MicroRNA family members mir-48, mir-84, and mir-241 function together to regulate developmental timing in Caenorhabditis elegans. Dev. Cell 9, 403–14.

5. Slack, F.J., Basson, M., Liu, Z., Ambros, V., Horvitz, H.R., and Ruvkun, G. (2000). The lin-41 RBCC Gene Acts in the C. elegans Heterochronic Pathway between the let-7 Regulatory RNA and the LIN-29 Transcription Factor. Mol. Cell 5, 659–669.

6. Reinhart, B.J., Slack, F.J., Basson, M., Pasquinelli, A.E., Bettinger, J.C., Rougvie, A.E., Horvitz, H.R., and Ruvkun, G. (2000). The 21-nucleotide let-7 RNA regulates developmental timing in Caenorhabditis elegans. Nature 403, 901–906.

7. Pepper, A.S.-R., McCane, J.E., Kemper, K., Yeung, D.A., Lee, R.C., Ambros, V., and Moss, E.G. (2004). The C. elegans heterochronic gene lin-46 affects developmental timing at two larval stages and encodes a relative of the scaffolding protein gephyrin. Development 131, 2049–2059.

8. Van Wynsberghe, P.M., Kai, Z.S., Massirer, K.B., Burton, V.H., Yeo, G.W., and Pasquinelli, A.E. (2011). LIN-28 co-transcriptionally binds primary let-7 to regulate miRNA maturation in Caenorhabditis elegans. Nat. Struct. & Mol. Biol. 18, 302.

9. Vadla, B., Kemper, K., Alaimo, J., Heine, C., and Moss, E.G. (2012). Lin-28 controls the succession of cell fate choices via two distinct activities. PLoS Genet. 8.

10. Moss, E.G., Lee, R.C., and Ambros, V. (1997). The cold shock domain protein LIN-28 controls developmental timing in C. elegans and is regulated by the lin-4 RNA. Cell 88, 637–646.

11. Miska, E.A., Alvarez-Saavedra, E., Abbott, A.L., Lau, N.C., Hellman, A.B., McGonagle, S.M., Bartel, D.P., Ambros, V.R., and Horvitz, H.R. (2007). Most Caenorhabditis elegans microRNAs are individually not essential for development or viability. PLoS Genet. 3, 2395–2403.

12. Ilbay, O., and Ambros, V. (2019). Pheromones and Nutritional Signals Regulate the Developmental Reliance on let-7 Family MicroRNAs in C. elegans. Curr. Biol. 29, 1735–1745.e4.

13. Karp, X., and Ambros, V. (2012). Dauer larva quiescence alters the circuitry of microRNA pathways regulating cell fate progression in C. elegans. Development 139, 2177–2186.

14. Bethke, A., Fielenbach, N., Wang, Z., Mangelsdorf, D.J., and Antebi, A. (2009). Nuclear hormone receptor regulation of microRNAs controls developmental progression. Science 324, 95–8.

15. Hammell, C.M., Karp, X., and Ambros, V. (2009). A feedback circuit involving let-7-family miRNAs and DAF-12 integrates environmental signals and developmental timing in Caenorhabditis elegans. Proc. Natl. Acad. Sci. U. S. A. 106, 18668–73.

16. Tabach, Y., Billi, A.C., Hayes, G.D., Newman, M.A., Zuk, O., Gabel, H., Kamath, R., Yacoby, K., Chapman, B., Garcia, S.M., et al. (2013). Identification of small RNA pathway genes using patterns of phylogenetic conservation and divergence. Nature 493, 694–698.

17. Ilbay, O., Nelson, C., and Ambros, V. (2019). C. elegans LIN-28 Controls Temporal Cellfate Progression by Regulating LIN-46 Expression via the 5’UTR of lin-46 mRNA. bioRxiv, 697490.

18. Fay, D.S., Stanley, H.M., Han, M., and Wood, W.B. (1999). A Caenorhabditis elegans homologue of hunchback is required for late stages of development but not early embryonic patterning. Dev. Biol. 205, 240–253.

19. Niwa, R., Hada, K., Moliyama, K., Ohniwa, R.L., Tan, Y.-M., Olsson-Carter, K., Chi, W., Reinke, V., and Slack, F.J. (2009). C. elegans sym-1 is a downstream target of the hunchback-like-1 developmental timing transcription factor. Cell Cycle 8, 4147–4154.

20. Abrahante, J.E., Daul, A.L., Li, M., Volk, M.L., Tennessen, J.M., Miller, E.A., and Rougvie, A.E. (2003). The Caenorhabditis elegans hunchback-like gene lin-57/hbl-1 controls developmental time and is regulated by microRNAs. Dev. Cell 4, 625–37.

21. Schwarz, G., Mendel, R.R., and Ribbe, M.W. (2009). Molybdenum cofactors, enzymes and pathways. Nature 460, 839.

22. Feng, G., Tintrup, H., Kirsch, J., Nichol, M.C., Kuhse, J., Betz, H., and Sanes, J.R. (1998). Dual Requirement for Gephyrin in Glycine Receptor Clustering and Molybdoenzyme Activity. Science (80-.). 282, 1321 LP – 1324.

23. Kneussel, M., Brandstätter, J.H., Laube, B., Stahl, S., Müller, U., and Betz, H. (1999). Loss of Postsynaptic GABA(A) Receptor Clustering in Gephyrin-Deficient Mice. J. Neurosci. 19, 9289 LP – 9297.

24. Fritschy, J.-M., Harvey, R.J., and Schwarz, G. (2008). Gephyrin: where do we stand, where do we go? Trends Neurosci. 31, 257–264.

25. Kirsch, J., Langosch, D., Prior, P., Littauer, U.Z., Schmitt, B., and Betz, H. (1991). The 93-kDa glycine receptor-associated protein binds to tubulin. J. Biol. Chem. 266, 22242–22245.

26. Fuhrmann, J.C., Kins, S., Rostaing, P., El Far, O., Kirsch, J., Sheng, M., Triller, A., Betz, H., and Kneussel, M. (2002). Gephyrin interacts with Dynein light chains 1 and 2, components of motor protein complexes. J. Neurosci. 22, 5393–5402.

27. Sabatini, D.M., Barrow, R.K., Blackshaw, S., Burnett, P.E., Lai, M.M., Field, M.E., Bahr, B.A., Kirsch, J., Betz, H., and Snyder, S.H. (1999). Interaction of RAFT1 with gephyrin required for rapamycin-sensitive signaling. Science 284, 1161–1164.

28. Antebi, A., Yeh, W.H., Tait, D., Hedgecock, E.M., and Riddle, D.L. (2000). daf-12 encodes a nuclear receptor that regulates the dauer diapause and developmental age in C. elegans. 1512–1527.

29. Ambros, V. (2004). The functions of animal microRNAs. Nature 431, 350–355.

30. Enright, A.J., John, B., Gaul, U., Tuschl, T., Sander, C., and Marks, D.S. (2004). MicroRNA targets in Drosophila. Genome Biol. 5, R1–R1.

31. John, B., Enright, A.J., Aravin, A., Tuschl, T., Sander, C., and Marks, D.S. (2004). Human MicroRNA Targets. PLOS Biol. 2, e363.

32. Bartel, D.P. (2004). MicroRNAs: Genomics, Biogenesis, Mechanism, and Function. Cell 116, 281–297.

33. Tsialikas, J., Romens, M.A., Abbott, A., and Moss, E.G. (2017). Stage-Specific Timing of the microRNA Regulation of lin-28 by the Heterochronic Gene lin-14 in Caenorhabditis elegans. Genetics 205, 251–262.

34. Nelson, C., and Ambros, V. (2019). Trans-splicing of the C. elegans let-7 primary transcript developmentally regulates let-7 microRNA biogenesis and let-7 family microRNA activity. Development, dev.172031.

35. Kim, H., Ishidate, T., Ghanta, K.S., Seth, M., Conte, D.J., Shirayama, M., and Mello, C.C. (2014). A co-CRISPR strategy for efficient genome editing in Caenorhabditis elegans. Genetics 197, 1069–1080.

